# Unraveling the effects of virtual reality overground walking on dynamic balance and postural control

**DOI:** 10.1101/2022.12.12.519831

**Authors:** Brian Horsak, Mark Simonlehner, Bernhard Dumphart, Tarique Siragy

**Affiliations:** Center for Digital Health & Social Innovation, St. Pölten University of Applied Sciences, St. Pölten, Austria; Institute of Health Sciences, Department of Health, St. Pölten University of Applied Sciences, St. Pölten, Austria

## Abstract

This study analyzed the effects of walking freely in Virtual Reality (VR) compared to walking in the real-world on dynamic balance and postural control. For this purpose nine male and twelve female healthy participants underwent standard 3D gait analysis while walking randomly in a real laboratory and in a room-scale overground VR environment resembling the real laboratory. The VR was delivered to participants by a head-mounted-display which was operated wirelessly and calibrated to the real-world. Dynamic balance was assessed with three outcomes: the Margin of Stability (MOS) in the anteroposterior (AP-MOS) and mediolateral (ML-MOS) directions at initial-contact, the relationship between the mediolateral Center of Mass (COM) position and acceleration at mid-stance with subsequent step width, and trunk kinematics during the entire gait cycle. We observed increased mediolateral (ML) trunk linear velocity variability, an increased coupling of the COM position and acceleration with subsequent step width, and a decrease in AP-MOS while walking in VR, but no change in ML-MOS when walking in VR. We conclude that walking in VR results in a less reliable optical flow, indicated by increased mediolateral trunk kinematic variability, which seems to be compensated by the participants by slightly reweighing sensorimotor input and thereby consciously tightening the coupling between the COM and foot placement to avoid a loss of balance. Our results are particularly valuable for future developers who want to use VR to support gait analysis and rehabilitation.

## Introduction

Augmented, Mixed, and Virtual Reality (VR) technologies extend our reality by merging the virtual with the real world thereby creating a fully immersive experience. The global market size for these technologies was approximately worth 31 billion U.S. dollars in 2021 and is projected to rise to almost 300 billion U.S. dollars by 2024 [1]. These developments raise great expectations for future applications in various domains such as entertainment, gaming, education, and fitness. However, the healthcare and diagnostics sector is expected to be most advanced by these technological developments. For example, VR has been and is still being intensively explored as a potential tool to aid gait rehabilitation for both clinical practice and research [2, 3].

Thus, it is not surprising that there is a constantly growing body of literature highlighting its potential for studying and aiding gait rehabilitation in patients with various movement or balance disabilities such as Parkinson’s Disease [4, 5], Cerebral Palsy [6], stroke [7–9], amputation [10], or aging [11–14]. While in the past VR has been traditionally used on treadmills, advancements in off-the-shelf head-mounted-display (HMD) technologies now allow to create immersive VR experiences where one can freely walk around and navigate through virtual overground environments whereby one’s real-life motion is reflected in the VR environment. Immersive overground VR environments offer partly unexplored possibilities for gait rehabilitation and research. They can be used to train activities of daily living [8], obstacle crossing [15], or allow the examination of effects for conditions which usually are difficult to evaluate such as visual impairments, dual tasks, or various environmental effects like diffuse lighting conditions or crowded places [16–18].

However, before such room-scale VR environments can be used as a purposeful tool in clinical and research practice it is important to understand if and to what extent that technology impacts our individual gait pattern. To date, there is only limited and partly inconsistent research available which specifically addressed this question for VR room-scale applications. Briefly summarized, researchers found a decrease in stride length [19], a decrease of walking speed, step length, and an increase of double support [20], while others reported a decrease in cadence [21]. In addition, Janeh et al. [22] reported that the decrease of walking speed during walking in the VR compared to the real-world did not normalize even during prolonged exposure to the VR environment. In contrast, one study found that a prolonged VR experience lead to a reduction in most VR-related gait adjustments [19]. The study of Yamagami et al. [23] is one of the few ones that assessed the effects of VR overground walking for people with Parkinson’s disease. They found similar effects, such as reduced walking speed, step length, and increased step width compared to the real-world laboratory. Interestingly, Martelli et al. [19] and Yamagami et al. [23] also reported a clear increase in variability for step width and length. In a previous study, our group examined the effects of a room-scale VR environment on healthy adults by means of standard clinical 3D full-body gait analysis [24]. The results of our study confirmed that individuals walk significantly slower, along with an increased double support time as well as increased variability in step width and foot off. Further, we found a markedly increased gait variability in the lower-extremity gait kinematic patterns. Changes in kinematic and kinetic gait patterns were small. Based on the currently available evidence it may be concluded that individuals seem to adapt towards a more conservative or cautious gait. Research demonstrates that individuals walking in challenging environments tend to adopt a more cautious gait pattern to mitigate the risk for falling [25–27]. This might also be the case for walking in VR room-scale environments. However, the simple analysis of spatiotemporal, kinematic, and kinetic gait patterns that the literature currently offers might not provide a complete answer to the question what the effects of VR are on dynamic balance and postural control.

For this reason a more specific analysis is needed to better understand potential effects of VR on our gait behavior. Gait is a dynamic motor task where the Center of Mass (COM) is held outside of the base of support (*BoS*) for the majority of the gait cycle, hereby defined as dynamic balance [28]. In this scenario, stability is achieved by the neuromuscular system counteracting the gravitational and joint reaction forces acting on the upper body to avert an uncontrolled fall [28]. One of the primary mechanisms to counteract these forces and maintain dynamic balance is through effective foot placement [28–30]. To achieve appropriate and effective foot placement, the neuromuscular system predicts the future kinematic state of the COM to determine upcoming foot placement at initial-contact [29]. In the anteroposterior direction, the passive dynamics of the body are exploited requiring minimal guiding information from automated subcortical and brain stem regions to determine foot placement [29, 30].

However, in the mediolateral direction, appropriate foot placement requires active sensory information processing from higher-level cortical regions [29]. Specifically, in this direction, the visual and vestibular systems provide information about head orientation which is then integrated with proprioceptive input from the trunk [29, 30]. The culmination of this process provides the neuromuscular system with an estimate of the dynamical state (position, velocity, and acceleration) of the COM which it then utilizes to determine upcoming mediolateral foot placement [29]. Indeed, previous research demonstrates that upcoming foot placement is determined by the kinematic state of the COM during the preceding contralateral mid-stance [29, 31–33]. As walking is a time-varying task with equally time-varying COM dynamical states, a constant sensory reweighing occurs to achieve foot placement [29]. However, it remains unexamined as to whether, and to what extent, VR affects this sensory motor reweighing as visual information is altered in the VR environment. Subsequently, this may impact determination of effective foot placement based on the movement of the COM which, in turn, may impact dynamic balance.

Therefore, this study attempts to analyze if walking through virtual room-scale environments affects our strategies in maintaining dynamic balance and postural control compared to walking in the real-world. For this purpose we revisited data from a recently published study [24]. That information could be of great value in assessing the potential fall risk individuals might be exposed to in VR room-scale environments. In addition, results will serve as a guide for future developers and researchers in creating more purposeful VR gait rehabilitation tools. Lastly, this research on healthy individuals should serve as a stepping stone to promote future studies evaluating if any potential fall risk exists for VR room-scale applications in various patient populations and aging demographics.

## Methods

### Participants and Study Design

This is a follow-up to a recently published study of our group on VR overground walking [24]. A convenience sample of 9 males and 12 female healthy volunteers (N = 21, age: 37.6 ± 8.6 yrs, weight: 70.8 ± 14.9 kg, height: 169.6 ± 6.8 cm) was recruited at our University’s campus located in the capital city of Lower Austria in Austria. All participants underwent standard 3D gait analysis in four randomly assigned walking conditions: the real laboratory (RLab, size: 11.9 × 5.4 m), a virtual laboratory resembling the real world (VRLab), and two differently sized versions of the VRLab. The latter two were not relevant for this study.

### Virtual Reality Room-Scale Environment

To immerse participants in the room-scale overground VR environment they wore a HTC Vive Pro HMD which was operated wirelessly and calibrated to the real-world. One HTC Vive 2.0 tracker was strapped to each foot to track and display the feet in VR in real-time (Fig 1). This allowed volunteers to have a visual indication of the current position in the VRLab and their body posture while navigating through the VR environment. Five HTC Vive Lighthouses (2.0) were used to continuously track the positions of the HMD and both trackers in the real-world.

**Fig 1.**
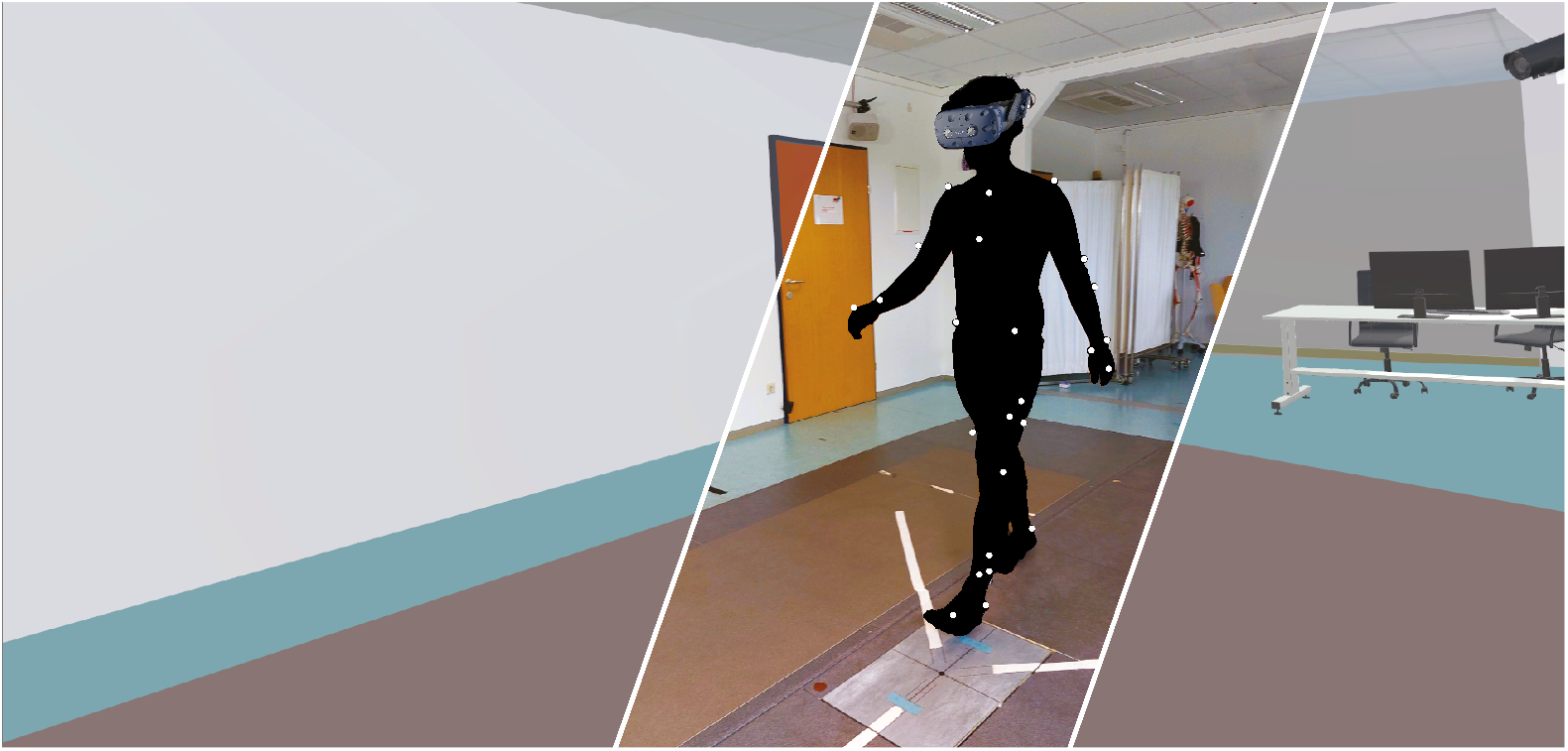
The real-world laboratory (RLab) partially superimposed with the virtual environment (VRLab) resembling the real-world. The Cleveland Clinic marker set was used for the lower body and the Vicon Plug-In-Gait model for the upper body. A wireless operated HTC Vive Pro was used to deliver the VR environment. HTC Vive 2.0 trackers were used to track the positions of the feet in real-time and display them in the VR. Note that we had to black out parts of the image to comply with the biorxiv policies.

### 3D Gait Analysis

A 12-camera motion capture system (Nexus, 2.11, Vicon, Oxford, UK) was used to collect trajectory data at 150Hz while one synchronized force plate (Kistler, Winterthur, CH) was used for collecting ground reaction force data at 300Hz. To collect participants’ kinematics, the extended Cleveland Clinic marker set [34] was used for the lower extremity and combined with the Vicon Plug-In-Gait model for the upper body. To determine the hip joint center, the regression equation from Davis et al. [35] was used. Kinetic data were filtered using a 4th order zero-lag butterworth filter with a cut-off frequency at 20Hz. All raw kinematic trajectories were filtered using the Vicon Nexus system integrated Woltring filtering routine with a MSA value of 15.

### Outcome Parameters

Data were analyzed in custom Python 3.8 scripts (using Spyder 4.1.5 within the Anaconda 3.0 suite). Gait events were determined via the force plate with a threshold of 20 N and by using an auto-correlation function and manual inspection for all steps before and after the force plate. All data were time normalized and expressed as a percentage of the gait cycle. To mitigate the influence of gait direction on our outcome parameters, i.e. walking not completely parallel to the anteroposterior axis of the laboratory coordinate system, a similar approach to [36] was chosen. Relevant trajectories for each stride were rotated by the angle determined between the gait direction and the anteroposterior direction of the laboratory for each stride separately. The rotation was then conducted about the pelvis center, defined as the mid-point between both hip joint center locations.

Dynamic balance and postural control was assessed with three outcomes: (i) the Margin of Stability (MOS) in the anteroposterior (AP-MOS) and mediolateral (ML-MOS) directions at initial-contact [37], (ii) the relationship between the mediolateral COM position and acceleration at mid-stance with subsequent step width [31], and (iii) trunk kinematics during the entire gait cycle.

The MOS was calculated bilaterally and defined as the distance of the extrapolated COM (*xCOM*) to the heel-marker for the AP-MOS or ML-MOS using equations 1 and See Fig 2 for a schematic description.

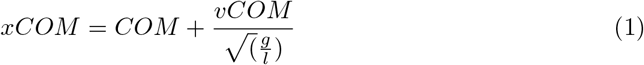

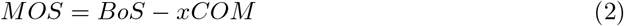

where *vCOM* is the velocity of the COM, *g* is the gravitational acceleration and *l* is the height of the inverted pendulum which was determined as the distance of the right/left heel marker to the COM at initial-contact. The position of the COM was estimated by using the cumulative mass and position of each anthropometric segment based on the Cleveland Clinic model [34] and Vicon’s Plug-In-Gait upper body model. Hof et al. [37] originally proposed to use the center of pressure as the *BoS*. However, we included all steps during walking along the walkway including those without force plate contact and therefore used the position of the heel-marker as an approximation of the COP during initial-contact. The MOS calculation is dependent on the foot and walking direction. To account for this we have used the (−1) term to correct for the directions of the *BoS* and *xCOM* vectors and ensure the MOS is positive when the *xCOM* is medial compared to the *BoS* or negative when the *xCOM* is anterior to the *BoS*. If the *xCOM* would be lateral to the *BoS* a person would have experienced a loss of balance. Note that, a negative AP-MOS indicates that the *xCOM* is progressing forward, which is the goal of forward walking. Since velocity is taken into account in the *xCOM*, the AP-MOS will lie outside the *BoS* during IC.

**Fig 2.**
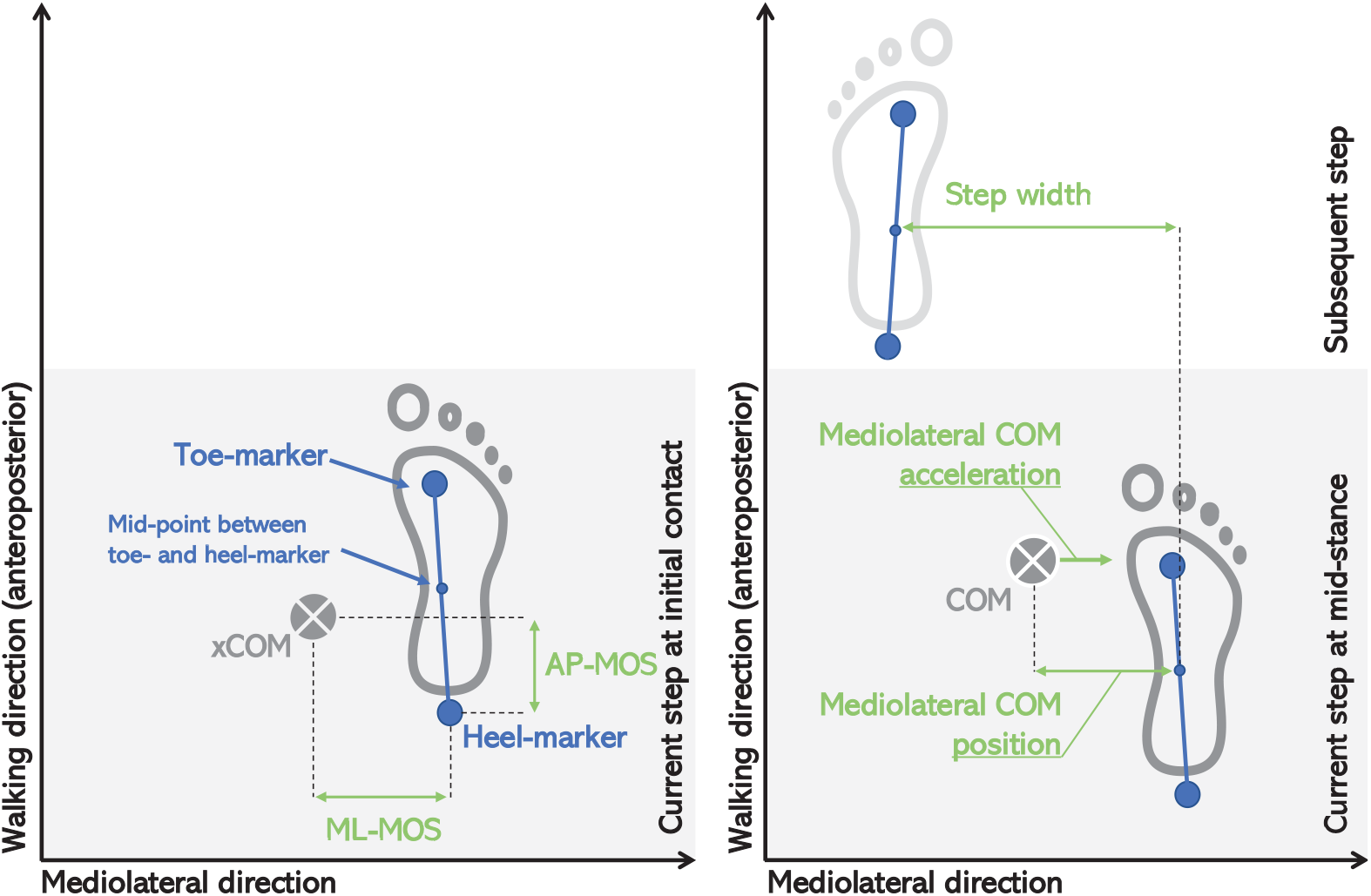
Schematic depicting some of our outcome variables relevant for this study. Left image: the Margin of Stability (MOS) was calculated for the mediolateral (ML-MOS) and anteroposterior (AP-ML) directions as the distance between the base of support (*BoS*) represented by the heel-marker at initial contact and the extrapolated center of mass position (*xCOM*). Right image: regarding the multiple linear regression, the mediolateral center of mass (*COM*) position and its acceleration during mid-stance were used to predict the subsequent step width. The *COM* position was defined as the horizontal distance between the vertical projection of the *COM* on the ground and the mid-point of the line connecting the heel- and toe-markers of the stance foot at mid-stance.

To assess the ability of the dynamical state of the trunk to determine mediolateral foot placement, we used similar to [31] a multiple linear regression model to relate the trunk COM position and its acceleration at mid-stance with the subsequent step width. Mid-stance was defined as the time point when the frontal velocity of the trunk COM was zero. The COM position was defined as the horizontal distance between the vertical projection of the COM on the ground and the mid-point of the line connecting the heel and toe markers of the stance foot at mid-stance. Step width was calculated as the frontal plane distance of the mid-point between toe- and heel-markers for both feet during two consecutive steps (see Fig 2). For the regression analysis, the trunk COM positions and accelerations at mid-stance for each left and right step were paired with the following step widths for each participant and lumped together in one sample. This resulted in a regression equation as followed for each walking condition. The explained variance in terms of the coefficient of determination (*R*^2^) served as outcome.

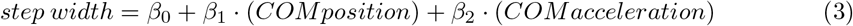

Lastly, to assess postural control we used the AP, ML, and vertical kinematics of the trunk to understand the impact of the VR on trunk kinematics. The T10-marker, placed right above the tenth thoracic vertebra, served as a surrogate to describe trunk motion. We have calculated the velocity profiles as well as the variability in terms of its standard deviation for each individual across the entire gait cycle and averaged them between the left and right steps to have robust and representative values per person.

### Statistical Analysis

Statistical analysis was conducted using SPSS Statistics 27 (IBM Corporation, NY, USA). Basic features of the data are reported as means and one standard deviation (SD), unless otherwise stated. Assumption of normality was checked by using a Shapiro-Wilk test and by inspecting the histogram of each variable.

As initial dependent pairwise tests did not indicate any significant asymmetries in MOS variables between the left and right side, values of both sides were averaged to have a single robust representation of each individual. Then, depending on the normality of the data either dependent t-tests or Wilcoxon signed rank tests were used to evaluate if a significant difference exists in the mean or median of the AP-MOS and ML-MOS for walking in RLab compared to walking in the VRLab. Due to the explorative character of this study and the small number of family-wise tests conducted we refrained from using Bonferroni-corrections to avoid high Type II errors. Alpha level was set to 0.05.

Regarding the assumptions for the regression analysis, a Durbin-Watson test was used to check for potential autocorrelation. Values in the range of 1 to 3 were interpreted as acceptable. Visual inspection of the histograms as well as skewness and kurtosis were used to test if residuals were approximately normal, the Goldfeld-Quandt test was used to test for homoscedasticity, and lastly the Variance Inflation Factor (VIF) was used to gauge multicollinearity between predictors. VIF values of greater than 2 were interpreted as a presence of moderate to strong multicollinearity.

Lastly, we utilized Statistical Parameter Mapping (SPM) and the SPM1D package (v.0.4.2) available for Python [38] to identify any differences in trunk kinematic velocity profiles and their variability between both walking conditions and separately for all three anatomical planes. For this purpose we used paired-sampled t-tests, or in case of none-normality of the data, non-parametric tests. Alpha level was set to 0.05.

## Results

A Shapiro-Wilk test indicated a normal distribution for the AP-MOS but a non-normal distribution for ML-MOS. A dependent t-test identified a significant difference in AP-MOS between walking in the RLab (−179 ± 34 mm) compared to walking in the VRLab (−158 ± 32 mm), t(20) = -4.991, ± p *<* .001. A Wilcoxon signed-rank test indicated no significant change for ML-MOS between walking in the RLab compared to the VRLab (Z = -.886, ± p = .375). Median ML-MOS was 19 mm and 20 mm for walking in RLab and VRLab, respectively. Results of the MOS are visualized in rain cloud box plots [39] in Fig 3.

**Fig 3.**
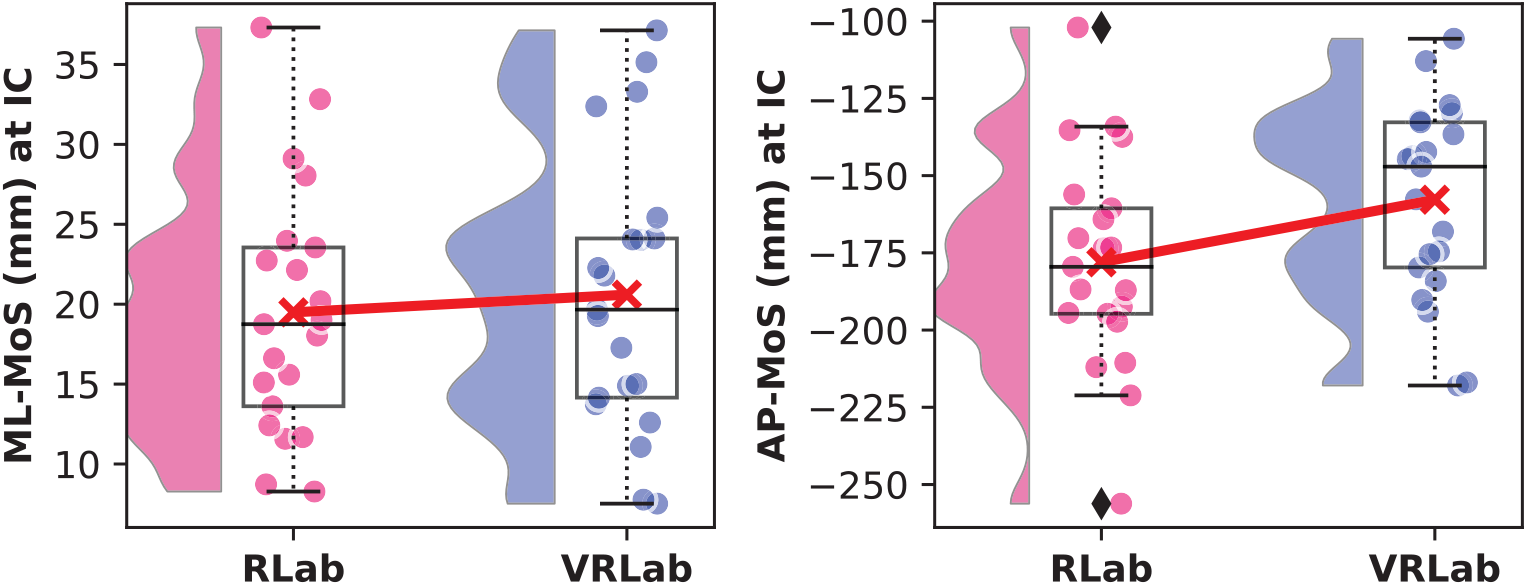
Mediolateral Margin of Stability (ML-MOS) and anteroposterior MOS (AP-MOS) at initial contact (IC) for walking in the real laboratory (RLab) compared to walking in the virtual laboratory resembling the real one (VRLab). The plot shows the data distribution (probability density function), the jittered raw data, the mean (red line), and a box plot showing quartiles where whiskers extend to the end of the data distribution except for outliers (diamonds).

Regarding the multiple regression both variables, mediolateral COM position and acceleration, significantly predicted step width and both variables added significantly to the prediction (p *<* 0.05) in both walking conditions. During walking in the RLab both variables accounted for 54% of the variance, F(2, 741) = 434.7, p *<* .001, *R*^2^ = .540. During walking in the VRLab 64% of the variance was explained, F(2, 812) = 717.4, p *<* .001, *R*^2^ = .639. Fig 4 shows a visualization of the 3D planes created by the regression equation for both walking conditions as well as the equation itself. From a goodness-of-fit perspective both regression models seemed appropriate and showed similar features in quality criteria. Durbin-Watson statistics was approximately 1.1, indicating moderate but acceptable auto-correlation, residuals were approximately normally distributed, the VIF (*<* 1.5) did not indicate high multicollinearity, and the Goldfeld-Quandt test was not significant (p *>* 0.6) indicating homoscedasticity.

**Fig 4.**
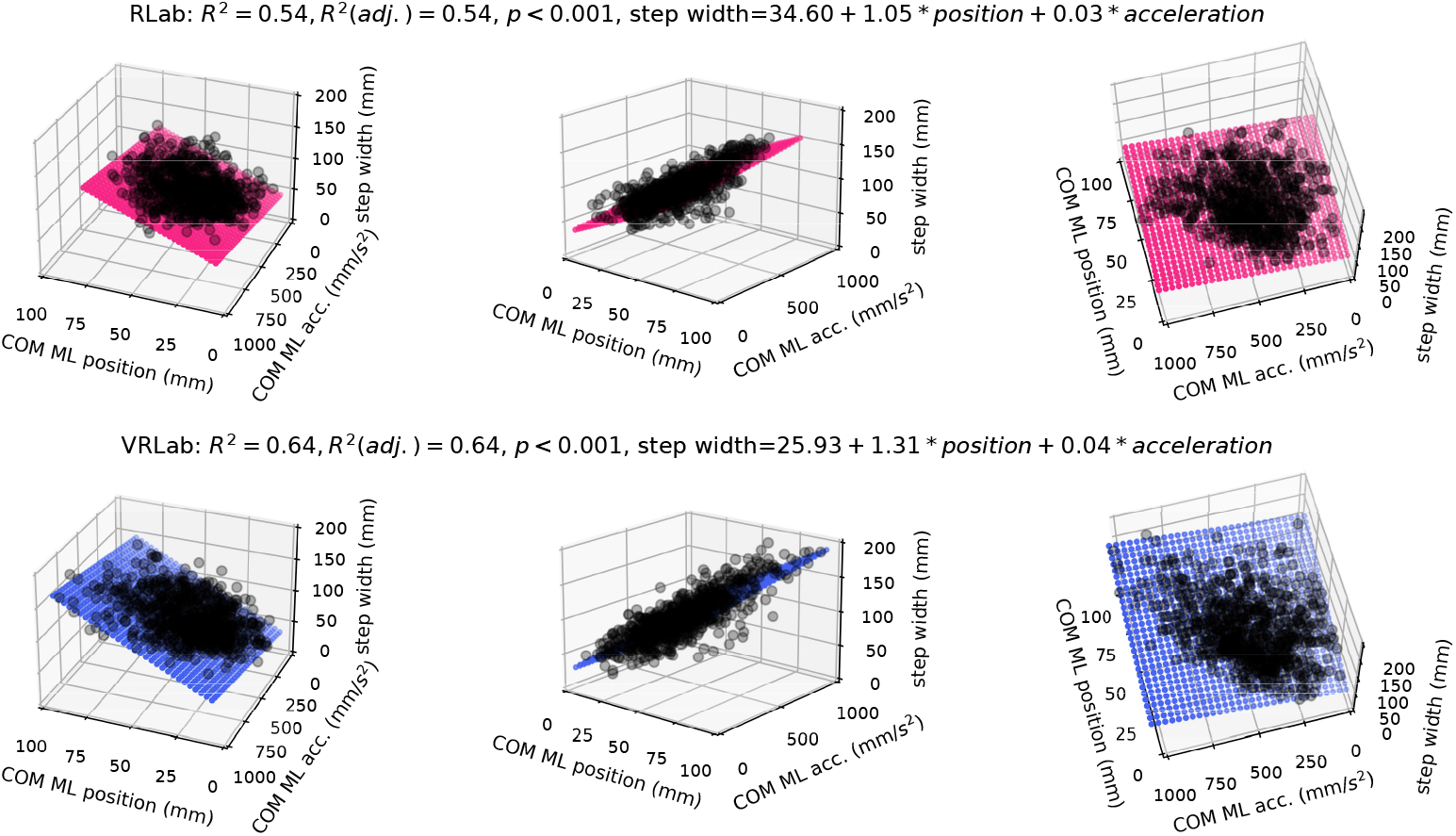
Three perspectives of a 3D visualization of the multiple linear regression model with mediolateral (ML) COM position (mm) and acceleration (mm/*s*^2^) as predictors for subsequent step width (mm). The upper row (pink) shows the model while walking in the real laboratory (RLab), the lower row (blue) depicts the model while walking in the virtual laboratory resembling the real one (VRLab).

SPM Shapiro-Wilk tests indicated non-normal distributions in some variables. Non-parametric paired sample SPM t-tests identified a highly significant decrease in absolute trunk velocity for the anteroposterior (−8.3 5.7%, ± p *<* 0.001) and vertical (−7.2 0.3%, ± p *<* 0.001) directions and a minimal increased mediolateral trunk velocity during early stance and early swing when walking in the VRLab compared to the RLab (see Fig 5). In addition, SPM identified an increased variability in mediolateral trunk velocity (+25.3 4.1%, ± p *<* 0.001) in the VRLab compared to RLab. No relevant changes in variability were found in the anteroposterior or vertical direction.

**Fig 5.**
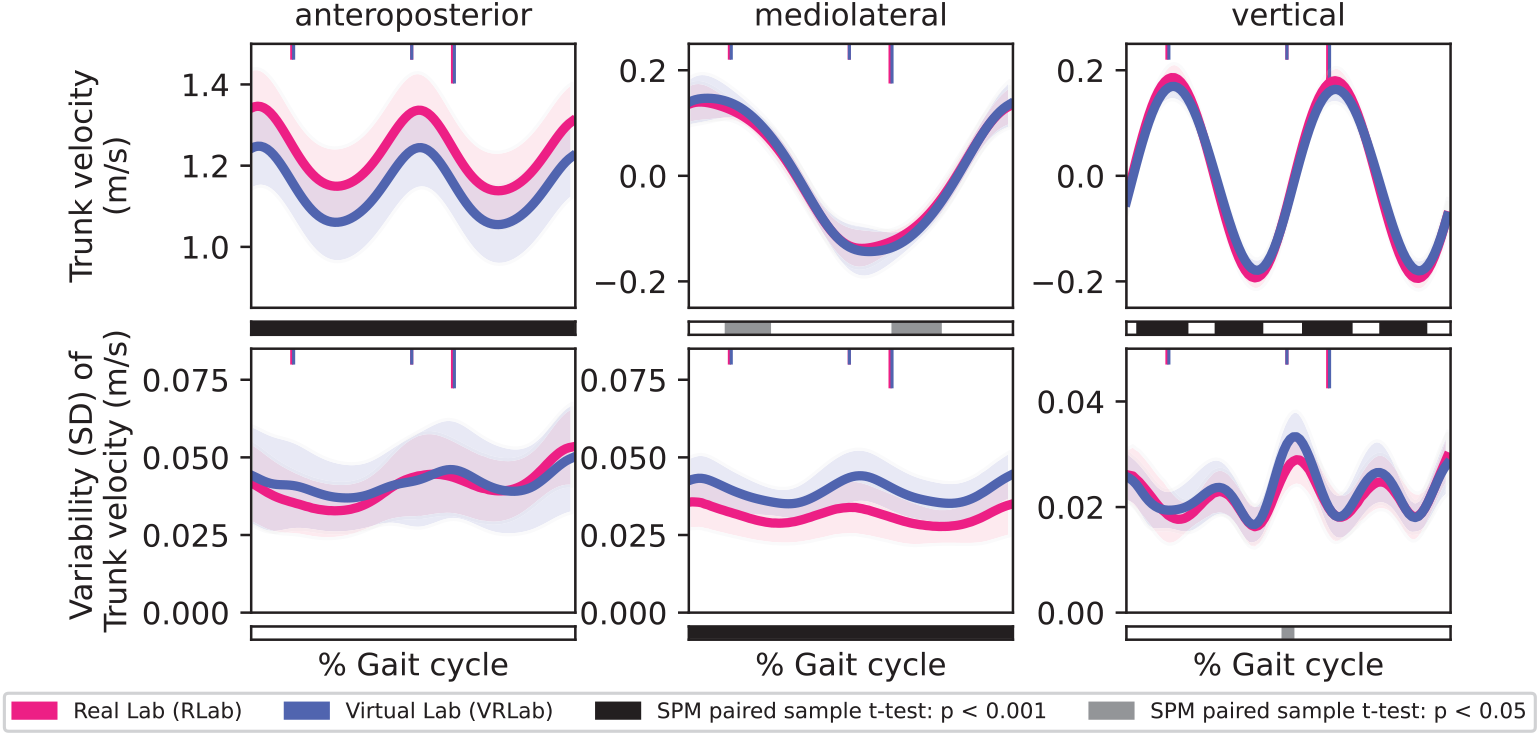
Kinematic profiles of the trunk velocity (first row) and its variability (second row) for the anteroposterior, mediolateral, and inferior-superior direction when walking in the real and the virtual laboratory. The bar below each graph indicates when the SPM {*t* } test statistic exceeded the critical threshold. Grey bars indicate a p *<* 0.05, black bars a p *<* 0.001.

## Discussion

This explorative study analyzed the effects of walking in a virtual overground room-scale environment with a head-mounted-display (HMD), compared to walking in the real-world, on dynamic balance and postural control in healthy adults. Our results demonstrated that walking in VR increases mediolateral trunk linear velocity variability, increases the relationship between the COM state and subsequent step width, decreases AP-MOS, but does not change the ML-MOS compared to walking in the real-world.

These findings are significant as instability and fall risk is greater in the mediolateral compared to the anterposterior direction while walking [28–30].

The increase in mediolateral trunk linear velocity variability may have stemmed from the difference in the visual scene between the VR and the real-world. Previous research demonstrates that postural control in the mediolateral direction, compared to the anteroposterior, requires the effective processing of visual information [29, 40].

Higher-level cortical structures integrate this input with somatosensory and vestibular input to provide information regarding the orientation and movement of the head relative to the trunk [29, 40, 41]. In turn, this provides the neuromuscular system with an estimation of COM kinematics during walking [29]. In our study, the increased mediolateral trunk velocity variability may indicate that participants had difficulty in effectively integrating the optical flow from the VR to control upper body posture. As a result, a sensorimotor perturbation may have arisen from the optical flow in the VR that destabilized our participants’ postural control. To rule out that the observed increased variability is only caused by the reduced walking speed, as slower walking speed is associated with increased kinematic trunk variability [42], we used a Pearson’s correlation to relate the change in walking speed (in terms of anteroposterior trunk velocity) and the increase in mediolateral variability of trunk velocity. No significant relationship was observed (see Figure 6) which further strengthens the notion that the altered optical flow in the virtual environment acted as a sensorimotor perturbation to our participants. Additionally, our finding is in line with current evidence which observed that mediolateral trunk kinematic variability increases during conditions of altered visual input [43]. Our results also partly underpin the results of Imaizumi et al. [44] who evaluated the effect of wearing a HMD on standing postural control by means of body sway analysis. Our results extend their findings to dynamic postural control. They found a largely increased body sway while wearing a HMD compared to not wearing one while with eyes opened. They assumed that the HMD changed the visual-feedback-based postural control by possibly reducing visual information about the position and motion of the head with respect to the surroundings thereby leading to a faster, more variable, and more adjustable body sway when wearing the HMD during standing.

**Fig 6.**
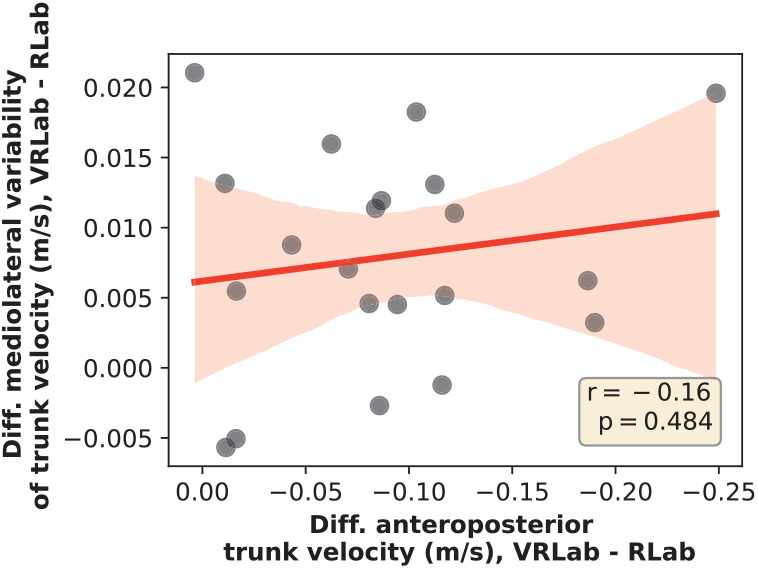
Linear Pearson’s correlation between the change in mediolateral variability of trunk velocity (y-axis) and the change in anteroposterior trunk velocity (x-axis) when walking in the virtual laboratory (VRLab) compared to the real laboratory (RLab). Differences were calculated as mean velocity during the entire gait cycle as VRLab - RLab.

Interestingly, despite the increases in mediolateral trunk variability, no changes occurred in the ML-MOS. Indeed, as the trunk accounts for the majority of our total-body mass, increases in trunk kinematic variability would disrupt the regular and sinusoidal movement of the COM during the gait cycle which threatens dynamic stability [30, 45]. However, in our preceding study [24] on this sample, we observed a simultaneous increase in participants’ step width variability. Although increased variability is a marker for falls in older adults [30, 46, 47], current evidence suggests that increases in spatiotemporal variability indicate foot placement adjustment to support a destabilized upper body in healthy young to middle-aged adults [30, 45]. Adjusting foot placement, as reflected by increased spatiotemporal variability, would account for the lack of findings in ML-MOS as participants modified their base of support to maintain their pre-existing level of dynamic stability in this direction. This is further strengthened by the fact that participants also displayed elements of a “cautious gait” strategy (slower walking speed, longer stride time, and increased double support time). When dynamic stability and postural control is threatened, individuals take slower steps to provide additional time for the COM to transition from the unloading to the loading leg during double-support [48, 49]. Additionally, the cautious gait strategy would account for the reduced AP-MOS observed during the VR condition compared to the real-world condition. Although vision is not closely linked to stability in the anteroposterior direction, [50] suggested that individuals implement this strategy to reduce the distance of the dynamical state of the COM to the base of support. As the dynamical state of the COM is located ahead of the base of support during steady-state walking, reducing the distance of the AP-MOS would facilitate the ability for individuals to return it within the base of support during a potential loss of balance such as when encountering an external perturbation. Thus, in our study, participants may have adopted a slower walking speed and a cautious gait strategy to reduce the likelihood of a balance loss in the event they encountered a potential perturbation in the real-world while wearing the HMD.

The ability to maintain a pre-existing level of mediolateral dynamic stability despite the altered optical flow in the VR may be due to sensorimotor reweighing which occurred. In a literature review, [29] discussed that during walking, multisensory information is integrated as a weighted average that is determined by the reliability of each individual sensory component. As such, when a specific component (vision) becomes less reliable, the sensorimotor system reweighs the afferent input to rely more heavily on alternative sensory information (somatosensory and vestibular). This reweighing may explain not only the lack of changes in ML-MOS, but also the increased *R*^2^ values during the VR condition. Indeed, we observed that COM kinematics (position and acceleration) accounted for 64 percent of the variance in step width during the VR condition compared to 54 percent during real-world walking. Hurt et al. [31] proposed that a stronger relationship between upper body kinematics and step width may indicate increased voluntary control to maintain dynamic balance while walking. As such, in our study, the potential sensorimotor reweighing that occurred, due to the altered optical flow in the VR, may have caused participants to consciously tighten the coupling between their COM and foot placement to avoid a loss of balance. However, it is unclear whether the change in magnitude of 10 percent, compared to a relative change of either 15 or 20 percent, holds a direct implication for dynamic balance.

Interestingly, Hurt et al. [31] reported differences of approximately 9% between young (24.5 ± 3.3 years) and older adults (60.6 ± 5.6 years). This might provide an approximate indication as older adults have reduced dynamic balance compared to healthy young adults [30]. [31], therefore, suggested this increase in the older adults was indicative of a more active strategy to control gait. However, they used a treadmill and the extent to which the relationship between variations in step width and COM state would be altered treadmill versus overground is presently not known. As such, future research should examine whether relative changes in magnitude affect an individual’s dynamic stability level as well as if treadmills further influence the relationship between step width and the COM state. Recent research indicates that VR based exergaming can effectively increase balance and reduce fear of falling in elderly [51–53]. VR is also increasingly used in patients with neurological disorders, such as Parkinson Disease. In a systematic review and meta-analysis, [54] recently confirmed that VR assisted balance training is highly effective in improving balance in patients with Parkinson’s Disease. While to date, the majority of research only uses non-immersive VR systems such as

Microsoft’s xBox and its wireless Kinect tracking system or non-immersive VR combined with treadmill training, the speed at which immersive VR technologies are currently developing suggests that there might be applications for fall-prevention and functional training in near future where highly immersive VR could play an important role. However, before such immersive VR technologies can serve as purposeful tools we need to fully understand the impact they have on our gait behavior and postural control. Unfortunately, as our study sample only comprised healthy individuals aged between 21 and 56 years, our results are limited in their generalizeability to the elderly population or patients with neurological disorders. The study by Yamagami et al. [23] is the only study we are aware of which evaluated the effect of VR on gait characteristics in patients with Parkinson Disease. They investigated whether Freezing-of-Gait (FoG) provoking VR environments exacerbate gait impairments associated with FoG compared to unobstructed VR and the physical laboratory. They found that walking speed was reduced and gait variability increased when people with Parkinson Disease walked overground in all VR environments. While both, their and our results point towards the same direction, there is still a need to better understand the impact of immersive VR on gait characteristics in various (patient) populations. This is important if we really want to fully exploit the potential of this rapidly advancing technology as a supportive tool for research and clinical care.

Lastly, our results need to be interpreted with caution when being used to inform the planning or development of prolonged exercise sessions using immersive VR as we do not know if the observed effects cease over a prolonged use or are independent by usage time and experience with VR. This is an important question which needs further attention.

## Conclusion

Recent studies in general observed a consistent pattern of gait adjustments when walking in VR overground environments compared to walking in reality [19–21, 23, 24]. Most frequently reported effects are reduced walking speed, increased gait variability, and step width which all point towards adjustments to a more cautious gait. Our study further underscores this idea, and is the first to provide an explanation from the perspective of dynamic stability and postural control. Our results indicate that the VR delivered to the participants with a HMD results in a altered optical flow, indicated by increased mediolateral trunk kinematic variability, which seems to be compensated by the participants by slightly reweighing sensorimotor input. Subsequently, participants consciously tightened the coupling between their COM and foot placement to maintain their already existing level of mediolateral dynamic stability. Although our results show some adjustments in dynamic stability and postural control, these should not be overestimated as we already showed that overall effects on the gait kinematic and kinetic patterns are rather small [24]. Immersive VR is a rapidly developing technology and it is reasonable to assume that VR and HMDs will become even more immersive in the near future thereby further reducing the effects they currently have on gait stability. Our study should thus be repeated in the foreseeable future with updated hardware and with various patient groups to support its application as a purposeful tool in healthcare.

## Ethics Statement

This study involving human participants was approved by the local ethics committee (GS1-EK-4/682-2020) and was performed in accordance with the relevant guidelines and regulations. All participants were informed prior to the study and gave written informed consent.

## Author Contributions

BH: Funding acquisition, Project administration, Conceptualization, Methodology, Writing - Original Draft, Formal analysis; MS: Methodology, Data Analysis, Data Curation, Review & Editing; BD: Methodology, Data Analysis, Data Curation, Review & Editing, TS: Data Analysis, Formal analysis, Writing - Review & Editing

## Acknowledgments

We would like to thank Lukas Richter for his thoughts and comments regarding the multiple linear regression analysis.

## Funding

This work received funding from the Austrian Research Promotion Agency (FFG) and the BMDW within the COIN-program (#866855) and from the Gesellschaft für Forschungsfö rderung NÖ (Research Promotion Agency of Lower Austria) within the Endowed Professorship for Applied Biomechanics and Rehabilitation Research

(SP19-004).

